# Machine Learning Informs RNA-Binding Chemical Space

**DOI:** 10.1101/2022.08.01.502065

**Authors:** Kamyar Yazdani, Deondre Jordan, Mo Yang, Christopher R. Fullenkamp, Timothy E. H. Allen, Rabia T. Khan, John S. Schneekloth

**Affiliations:** Chemical Biology Laboratory, Center for Cancer Research, National Cancer Institute, Frederick, MD 21702-1201, USA; Ladder Therapeutics, USA

## Abstract

Small molecule targeting of RNA has emerged as a new frontier in medicinal chemistry, but compared to the protein targeting literature our understanding of chemical matter that binds to RNA is limited. In this study, we report <u>R</u>epository <u>O</u>f <u>BI</u>nders to <u>N</u>ucleic acids (ROBIN), a new library of nucleic acid binders identified by small molecule microarray (SMM) screening. The complete results of 36 individual nucleic acid SMM screens against a library of 24,572 small molecules are reported (including a total of 1,627,072 interactions assayed). A set of 2,003 RNA-binding small molecules is identified, representing the largest fully public, experimentally derived library of its kind to date. Machine learning is used to develop highly predictive and interpretable models to characterize RNA-binding molecules. This work demonstrates that machine learning algorithms applied to experimentally derived sets of RNA binders are a powerful method to inform RNA-targeted chemical space.

## Introduction

Recent years have seen an increased interest in targeting RNA with small molecules as an approach to develop novel chemical probes and therapeutics.^[1]^ The potential for developing transformative therapeutics that target RNA is considerable. Of the ∼3 billion base pairs in the human genome, estimates suggest that nearly 85% of this information is transcribed into RNA.^[2]^ In contrast, just ∼1.5% of transcribed sequences code for polypeptides.^[3]^ Research into the non-coding functions of RNA has revealed numerous disease-relevant regulatory mechanisms that could conceivably be modulated by RNA-targeting molecules.^[1a, 4]^ While estimates indicate that less than 15% of the proteome is “druggable”,^[5]^ a far greater proportion of proteins have been implicated in disease states, and targeting the mRNA of “undruggable” proteins could represent a unique approach to impact the expression of pathogenic proteins.

To date, antisense molecules such as siRNAs, shRNAs, PNAs, and LNAs have been widely used to target RNA.^[6]^ However, antisense technologies often bring challenges with distribution or cell permeability,^[7]^ making small molecules that target RNA a highly attractive option. Early efforts to target RNAs with small molecules focused on molecules that recognize grooves in helices or non-specific chemical scaffolds such as aminoglycosides.^[8]^ However, because RNA is single-stranded, it can form intramolecular base pairs and fold into a diverse array of highly complex structures that are capable of forming hydrophobic pockets that are likely to accept small molecule ligands.^[9]^ Structures in RNA can be functionally relevant,^[10]^ and may therefore represent opportunities to perturb RNA with small molecules.^[11]^ To date, viral RNAs, riboswitches, ribosomal RNA, microRNAs (miRNAs), long non-coding RNAs (lncRNAs), and mRNAs have been targeted with small molecules.^[1a, 1b, 8, 12]^ Compared to protein targets, RNA poses unique challenges due to its anionic phosphodiester backbone, the highly dynamic nature of many reported RNA structures, and a relative lack of atomic resolution structures of RNA. More work is needed to understand the chemical properties that drive RNA recognition by small molecules, and what makes a good RNA target.

While numerous studies have detailed the chemical properties of protein-binding molecules and what factors are likely to lead to the successful development of an inhibitor,^[13]^ a comparably small number of studies have broadly described the properties of RNA-binding molecules. Efforts by the Hargrove laboratory,^[14]^ Disney laboratory,^[15]^ and a Merck research group^[16]^ have described a number of chemotypes and chemical properties enriched in RNA binding small molecules. In addition, several databases exist to catalog RNA- and DNA-binding molecules. In this area, notable contributions include INFORNA (based on 2-dimensional combinatorial screening)^[15c]^, R-BIND^[14d, 14e]^ (focused on biologically active RNA-binding molecules) and NALDB^[17]^ (which catalogs DNA- and RNA-binding molecules), all of which represent important resources for understanding chemical matter that interacts with nucleic acids.^[14e, 17]^ However, to date no laboratory has published full results from high-throughput screening against diverse RNA targets (including both hits and complete library composition). Such data would be uniquely valuable for developing advanced algorithms to characterize and understand RNA-binding chemical space.

Here, we disclose the complete results of 36 small molecule microarray (SMM) screens against nucleic acid targets. These targets represent diverse RNA structures and several DNA G-quadruplexes screened against a library of 24,572 commercially available drug-like small molecules. This dataset includes replicate assays and some oligos screened against a subset of the library, for a total of 1,627,072 interactions probed. Comparative analysis of these screens reveals information about the targetability of classes of nucleic acid structure and frequency of selective hit identification. While DNA structures are included in the dataset, much of the analysis herein is focused on RNA targets. We observe that many RNA-binding small molecules fall within “drug-like” chemical space, as defined by traditional medicinal chemistry parameters. However, such parameters were insufficient to distinguish protein and RNA binders in this dataset. We therefore enumerated ∼1,600 chemical properties of our libraries and analyzed experimentally identified RNA-binding molecules using machine learning approaches. Logistic regression and neural network models point to a complex interplay of dozens of molecular descriptors able to distinguish between RNA- and protein-binding compounds. This work reveals that despite RNA and protein binders having similar drug-like properties, molecular determinants related to nitrogenous and aromatic rings, Van der Waals surface area, and topological charge can help distinguish between molecules of these two sets. This work defines a boundary between RNA- and protein-binding small molecules that will facilitate the design of chemical libraries as well as individual ligands that target RNA structures.

## RESULTS

### Composition of the ROBIN Library

To screen a collection of drug-like small molecules against nucleic acid targets, we assembled a library of commercially available compounds from diverse chemical sources that were compatible with SMM screening (each compound contained an amine or alcohol for attachment to the SMM slide). A custom set of 11,515 compounds was obtained from ChemDiv, 9,595 compounds from ChemBridge, and 1,593 compounds from the National Cancer Institute Diversity Set. In addition, 1,869 compounds were acquired from the National Center for Advancing Translational Sciences (NCATS) Mechanism Interrogation PlatE (MIPE) library,^[18]^ representing mechanistically characterized, biologically active drugs and probes. In general, compounds from commercial vendors were chosen based on modern medicinal chemistry parameters such as molecular weight, hydrogen bond donors/acceptors, polar surface area, predicted solubility, and filtered to generally avoid most reactive chemical groups or assay interference chemotypes (e.g., “PAINS”^[19]^). A full description of the chemical library including SDF files and the analysis of chemical composition, are included in Supporting Information. This library was screened against 27 RNA and 9 DNA targets belonging to diverse structural classes such as hairpins, G-quadruplexes (G4), pseudoknots (PK), three-way junctions (TWJ), and triple helices (TH). The results of several individual screens using this library have been published previously by our laboratory.^[11, 20]^ We used the results of SMM screens for 36 nucleic acid targets against this library to compile the ROBIN library, a collection of 2,188 unique small molecules that were a hit for at least one nucleic acid target. A description of each nucleic acid target screened is described in Table S1. The protocol used in our SMM screens and our criteria for hit identification have been previously described.^[21]^ Summarized data for these SMM screens including hit rates for individual structural classes of nucleic acids are reported in Table 1. The mean hit rates for all structural classes were less than one percent, comparable to what is seen for many protein targets in unbiased screening efforts. Selective hit rates, reporting only compounds that bound to the target of interest and no other target, were considerably lower. The hit rate, structure class, and sequence of each individual nucleic acid target can be found in Table S2.

**Table 1.** Statistics for hit rates of 36 SMM screens classified by nucleic acid structure type. Included are hit rate (average hit rate for oligonucleotide targets in that class of structure) and selective hit rate (average hit rate for oligos, considering only compounds that bind to that target and no others).

| Structure Type | Count | Hit Rate (%) | Selective Hit Rate (%) |
| --- | --- | --- | --- |
| DNA G4 | 9 | 0.45 ± 0.18 | 0.12 ± 0.15 |
| RNA G4 | 6 | 0.78 ± 0.26 | 0.22 ± 0.15 |
| RNA Hairpin | 14 | 0.48 ± 0.26 | 0.16 ± 0.12 |
| RNA Pseudoknot | 3 | 0.85 ± 0.15 | 0.33 ± 0.07 |
| RNA 3WJ | 2 | 0.79 ± 0.18 | 0.23 ± 0.17 |
| RNA Triple Helix | 2 | 0.61 ± 0.23 | 0.21 ± 0.02 |

To quantitatively describe selectivity for hit molecules, we calculated Gini coefficients for each of the 2,003 RNA-binding compounds. The Gini coefficient is a metric derived from economic research on inequality that has been proposed as a way to quantitatively measure the selectivity of chemical probes including for RNA binders.^[22]^ For example, a molecule that binds to only one target out of many would have a theoretical Gini coefficient of 1 and a molecule that binds to all targets equally would have a Gini coefficient of 0. A value above 0.75 has been proposed as a reasonable cutoff for selective chemical probes.^[23]^ In our dataset, 1,287 compounds had Gini coefficients above 0.75, while 716 were below 0.75 (Supporting Information, Figure S1). Thus, 36% of the identified hits displayed promiscuous binding to RNA, and selective hits were identified for each structure screened.

### Comparison of ROBIN RNA-Binding Library with FDA-approved Drugs

While compounds like aminoglycosides are capable of binding to nucleic acids with high affinity, the physical properties and lack of selectivity of these molecules make them poor candidates for further development as targeted therapeutics. To examine whether ROBIN RNA binders (2,003 compounds) have properties that more closely resemble those of classically defined “drug-like” molecules, we compared these compounds to FDA-approved drugs in a commercially available library from Selleck Chemicals. The library of FDA-approved drugs was filtered before analysis to remove nucleic acid derivatives, molecules with a high molecular weight (> 1000 Da), and molecules containing platinum or other metals for a final set of 2,350 compounds. We generated kernel density estimate (KDE) plots for six traditional medicinal chemistry parameters as described by Lipinski^[24]^ and others^[25]^ (molecular weight (MW), number of hydrogen bond acceptors (HBA) and donors (HBD), number of nitrogens, total polar surface area (TPSA), and Wildman-Crippen LogP (SLogP)). These distributions are shown for the FDA-approved set, ROBIN RNA-Binding library, and the remaining compounds in SMM screening that did not bind to any RNA (Figure 1A).

**Figure 1.**
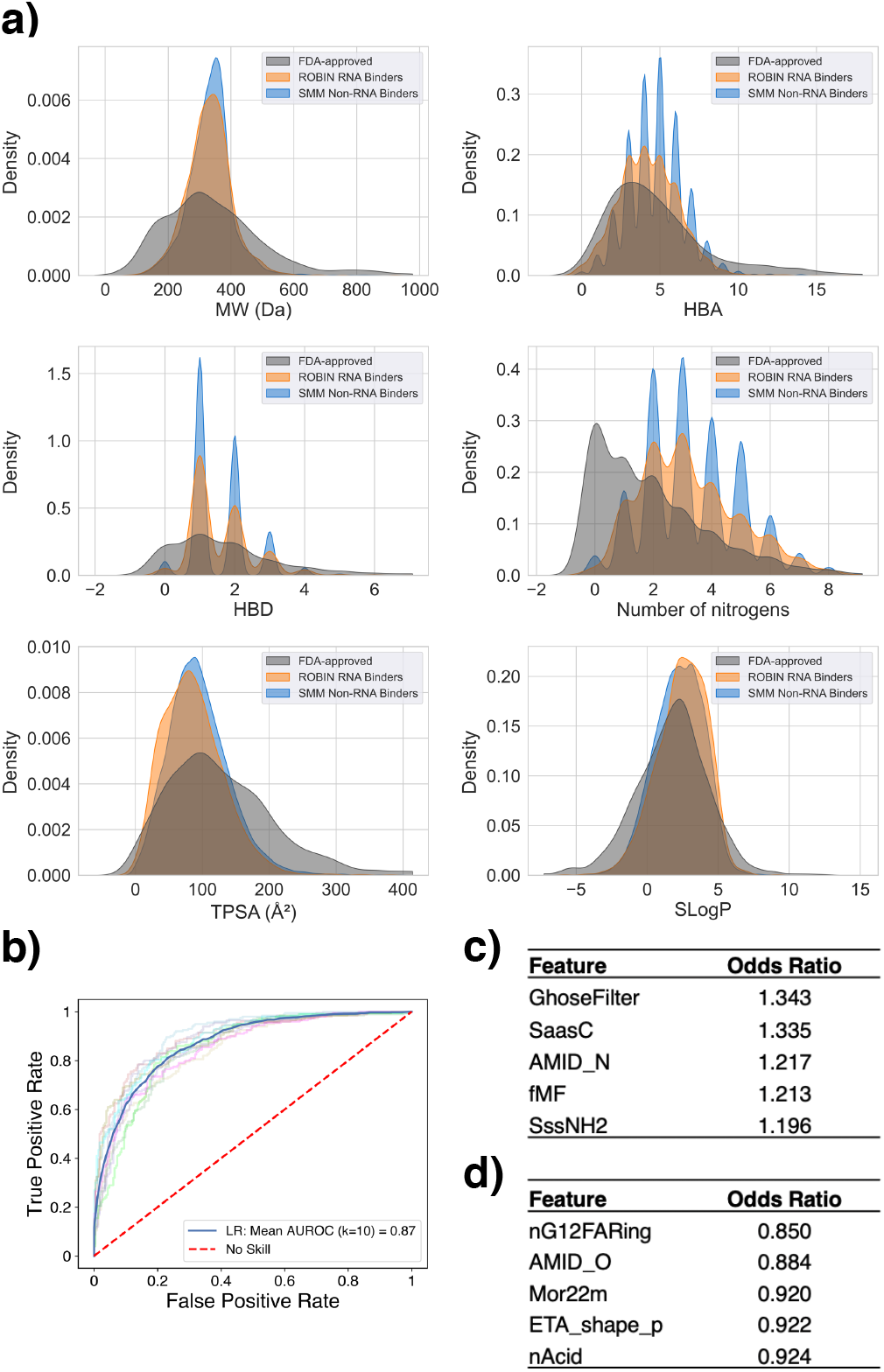
Comparison of ROBIN RNA binders and FDA-approved drugs. **A)** Kernel Density Estimate (KDE) plots of distributions of six common medicinal chemistry parameters for FDA-approved drugs (grey), ROBIN RNA binders (orange), and compounds from the SMM library that did not score as hits for any RNA (SMM Non-RNA Binders, blue). **B)** Receiver Operating Characteristic (ROC) curve for classification of ROBIN RNA binders and FDA-approved drugs using LASSO logistic regression. **C)** Five features with the highest odds ratios identified by LASSO. **D)** Five features with the lowest odds ratios identified by LASSO.

We used the Mordred software package^[26]^ to generate 1,827 two-dimensional and three-dimensional molecular descriptors for each compound in ROBIN and the FDA-approved set, of which 1,664 features were successfully enumerated. The Mordred package was chosen because it is widely utilized, freely available, and high-performance (capable of parallel computation). We used the generated Mordred features to inform a logistic least absolute shrinkage and selection operator (LASSO)^[27]^ regression model to perform binary classification on these compound sets. LASSO logistic regression is a penalized regression model that performs feature selection and removes redundant covariates automatically. The main advantage of LASSO lies in its ability to arrive at sparse solutions due to applying a L1-norm penalty on its coefficients. This analysis arrived at a panel of 41 molecular descriptors and was able to achieve an outstanding classification performance with mean area under the receiver operating characteristic (AUROC) of 0.87 using 10-fold cross-validation (Figure 1B). Figures 1C and 1D illustrate features with the five highest and lowest odds ratios, respectively. KDE plots showing the distribution of these 10 features are also shown for the FDA-approved set and the ROBIN RNA-binding library in the Supporting Information.

### Classification of RNA-binding and Protein-binding Compounds

To visualize the distribution of ROBIN RNA binders within chemical space, we utilized a recently developed method known as TMAP (Tree MAP)^[28]^. TMAP enables visualization of large high-dimensional datasets in a human interpretable manner using a two-dimensional minimum spanning tree (MST). Tree-like configuration of this dimensionality reduction technique attempts to preserve the global and local patterns of the dataset and can outperform related algorithms like t-SNE and UMAP in performance. TMAP utilizes this tree-like layout to indicate clusters as branches. To illustrate the distribution of RNA binders in chemical space, we assembled 2,350 FDA-approved drugs as described earlier and 10,000 randomly selected compounds from BindingDB^[29]^ (which catalogs small molecules that target proteins) with *K*_*d*_ or *K*_*i*_ < 10 nM, indicating tight binding to protein targets. We also added 2,003 compounds from the ROBIN RNA-binding library as representatives of RNA-binding chemical space. We encoded these chemical libraries with extended connectivity fingerprint up to 4 bonds (ECFP4) fingerprints that have been widely used in drug discovery studies^[30]^. Figure 2A illustrates the TMAP for these three datasets. In addition, Figure 2A illustrates a detail of the TMAP, where related structures are found in all three sets.

**Figure 2.**
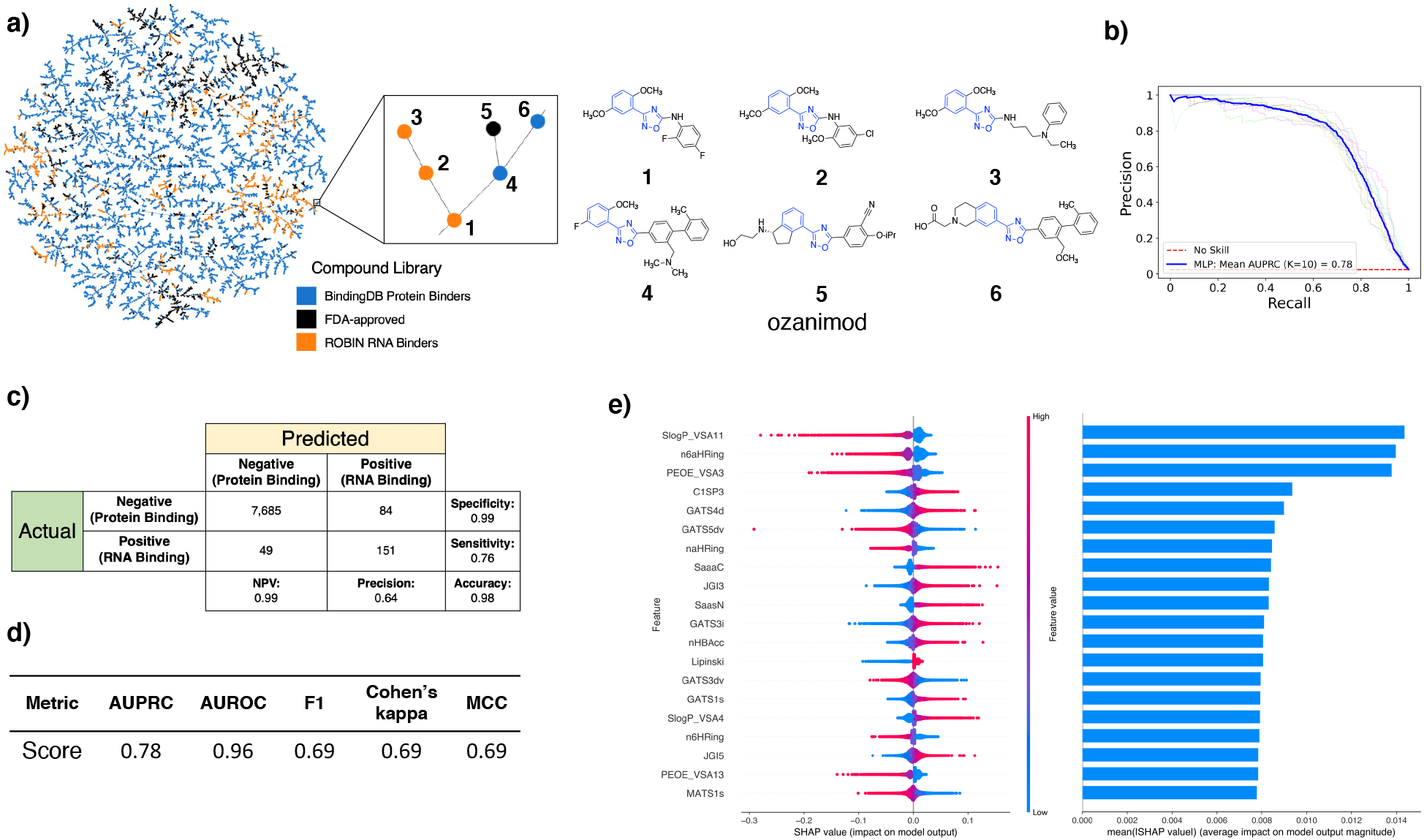
Classification of ROBIN RNA binders and protein binders. **A)** Left, TMAP of 10,000 BindingDB protein binders (blue), 2,350 FDA-approved drugs (black), and 2,003 ROBIN RNA binders (orange) encoded with ECFP4 fingerprints. Right, six structures shown from a detail of the TMAP illustrating related molecules on a branch. **B)** Precision recall (PR) curve for classification of augmented ROBIN RNA binders and BindingDB protein binders using the multi-layer perceptron (MLP) model. **C)** Confusion matrix showing the performance of the MLP on the test/holdout set. NPV refers to the negative predictive value. **D)** Performance metrics for the MLP model on the test/holdout set. **E)** Left, beeswarm plot illustrating how the top 20 most important features in the MLP impact the model’s output. Right, bar chart showing mean absolute SHAP values for each feature in the beeswarm plot. Each row of the bar chart is aligned with feature rows of the beeswarm plot.

To compare ROBIN RNA binders to protein-binding small molecules, we extracted the entire set of compounds from BindingDB with reported *K*_*d*_ or *K*_*i*_ < 10 nM to a protein target (77,678 compounds). We used the Mordred chemical descriptors to classify this set from the library of 2,003 ROBIN RNA binders using a class-weighted LASSO logistic regression model. Our classification performed with a mean area under the precision-recall curve (AUPRC) of 0.38 using 10-fold cross-validation (see Supporting Information). Precision-recall (PR) curve is used in this case as receiver operating characteristic (ROC) curves can be misleading in cases of high class imbalance. This model, however, did not generalize to several experimentally validated compounds from the literature, potentially due to biases introduced by the minority class (ROBIN RNA binders) being outnumbered 39 to 1 in the training set. To generate a more class-balanced training set, we used an oversampling strategy to augment ROBIN RNA binders. Before augmenting the data, we removed 10% of the RNA and protein binders for an unbiased test/holdout set. Next, we split the remaining compounds in the experimental sets (1,803 RNA binders and 69,909 protein binders) into 10 stratified folds for cross-validation. Experimental RNA binders in each training set of the cross-validation (1,803 compounds) were augmented 30 fold by identifying related structures in the ZINC^[31]^ database. Analogs were selected to have Tanimoto similarity > 0.85^[32]^, molecular weight > 250, and cLogP between -1 and 5. If insufficient analogs were identified, the RNA binder was replicated until a total of 30 compounds were reached for each case. We trained the LASSO logistic regression model on this augmented training set, and each round of cross-validation was tested against the non-augmented set of RNA and protein binders (180 and 6,991 compounds, respectively). With the augmented ROBIN dataset, the LASSO logistic regression algorithm achieved mean AUPRC of 0.37 with 10-fold cross-validation. This model identified 294 molecular descriptors with non-zero coefficients (see Supporting Information for more details on this model). Notably, due to a large number of multicollinear molecular descriptors within the dataset, non-zero coefficients of these do not imply that these features are uniquely more important than other collinear molecular descriptors. Rather, LASSO randomly omits redundant covariates to arrive at a sparse solution. The selection of these 294 molecular descriptors by LASSO indicates that they are sufficient, but not necessarily unique, in successful binary classification of RNA- and protein-binding small molecules.

In addition to LASSO logistic regression, we utilized a more complex non-linear model to achieve a superior classification performance. We used a class of feedforward neural networks known as a multilayer perceptron (MLP). Using the same augmentation and cross-validation strategy, we reached a mean AUPRC score of 0.78 (Figure 2B) on our augmented set. A confusion matrix and other relevant performance metrics are shown in Figures 2C and 2D, respectively. The architecture of the MLP is shown in the Supporting Information. Despite the outstanding performance of the neural network, this algorithm is a “black box” algorithm meaning it is difficult to interpret how the algorithm handles the chemical features in its hidden layers.

Over the past several years, there have been significant advances in developing algorithms capable of calculating feature importance in predictions of complex models such as neural networks. One such method is SHapley Additive exPlanations (SHAP).^[33]^ SHAP draws its inspiration from coalition game theory and provides local explanations for the importance of features in individual predictions. SHAP provides a higher payout to features that contribute more to the model performance. Higher absolute SHAP values for a chemical descriptor would therefore indicate higher importance of that feature to the model. In our SHAP analyses, protein binders are labeled as 0 and RNA binders as 1. Thus, higher SHAP value of a feature for a compound pulls the compound more toward RNA binding. In contrast, features with lower SHAP values push the compound more toward protein binding. Applying SHAP analysis to our MLP model, we identified the top 20 features with the highest mean absolute SHAP values. A beeswarm plot of these chemical features showing how they impact the model is illustrated in Figure 2E. Each row in the beeswarm plot corresponds to a feature and all compounds in the dataset are plotted for each feature as dots. The position of each dot along the X-axis denotes the SHAP value of the compound for that feature. In addition, the dot color indicates the feature value for each compound. For example, higher 6-membered aromatic hetero ring count (n6aHRing) values are associated with lower SHAP values meaning higher n6aHRing values pull the compound more to the protein binding side. It is clear that a complex set of chemical features relating to properties such as Van der Waals surface area (e.g., SlogP_VSA11, PEOE_VSA3, SlogP_VSA4, PEOE_VSA13), topological charge (e.g., JGI3, JGI5), aromaticity and nitrogen ring systems (e.g., n6aHRing, naHRing, SaasN, SaaaC), SP^3^ character (C1SP3), and hydrogen bond acceptors (nHBAcc) can be predictive for RNA binding within a set of drug-like chemical scaffolds.

We then proceeded to test the performance of our MLP model on several compounds previously demonstrated to bind to RNAs or proteins but not included in our datasets. These compounds include ADQ (a small molecule binder of HOTAIR),^[34]^ a compound that binds to the HIV transactivation response (TAR) hairpin (compound 4), _[35]_ ribocil-A (a synthetic ligand for the FMN riboswitch),^[36]^ and tetracycline (binds to the 30S and 50S subunit of bacterial ribosomes) (Figure 3A).^[11]^ We also selected four widely used protein-targeting FDA-approved drugs with well-validated mechanisms of action including imatinib, ibrutinib, lovastatin, and nevirapine (Figure 3B). Notably, these four drugs are included in our SMM library but were not a hit for any RNA target. The neural network’s performance on these eight examples is illustrated in Figure 3. Here, it is clear that molecules previously validated to bind to RNA are predicted to have a high chance of RNA binding. In contrast, known protein binders not observed to bind to RNA in SMM screening are scored far lower in the model even though they were not included in the ML training.

**Figure 3.**
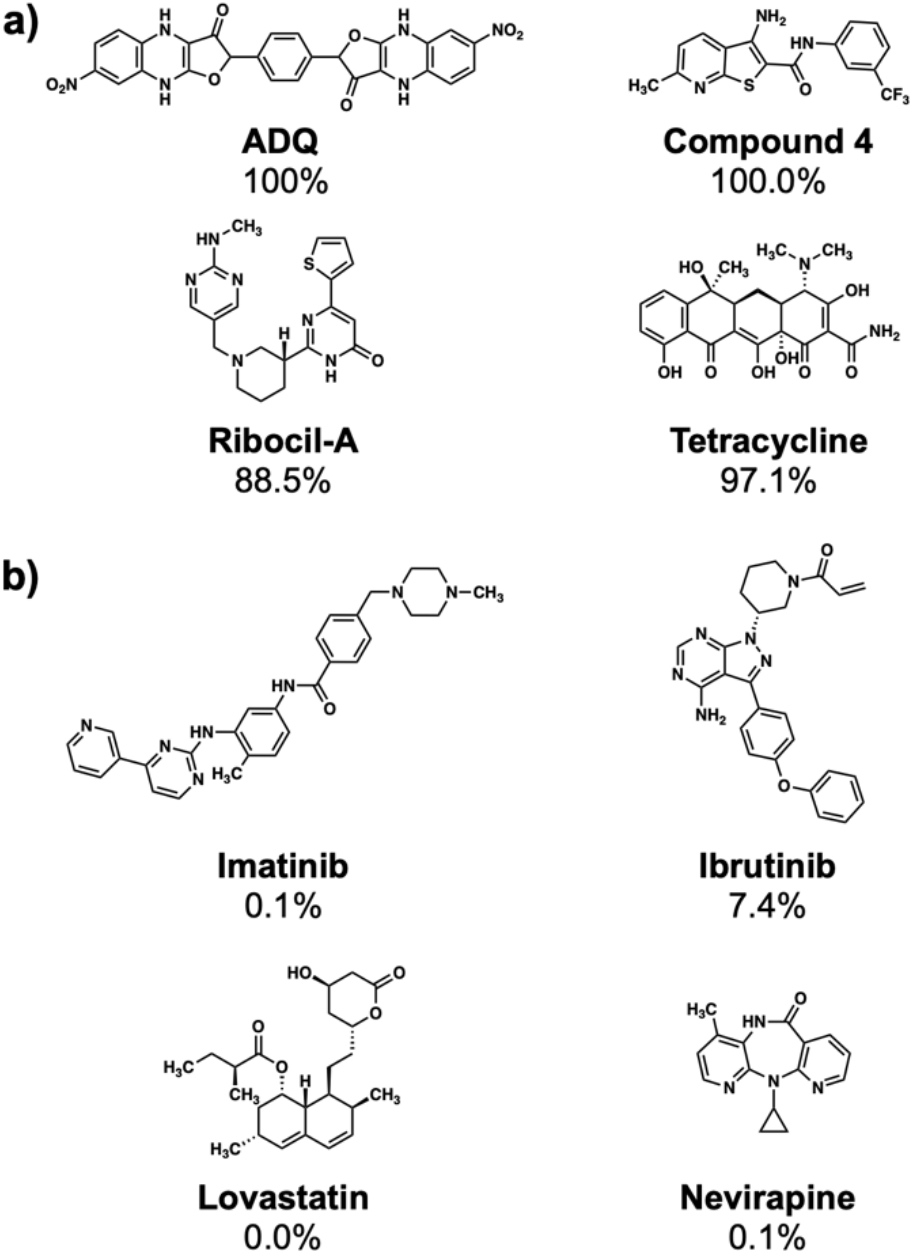
Performance of the MLP model on selected known RNA and protein binders. **A)** Model performance on four known RNA binders not included in the SMM screening library. **B)** Model performance on four known protein binders. Note that all the protein binders were printed on SMMs and showed low/no binding to RNA targets screened. In each case, the value reported represents probability of RNA binding relative to protein binding as predicted by the MLP model.

### Substructure Searching

To identify substructures enriched in selective RNA binders, we curated a set of 423 functional groups and ring structures commonly found in FDA-approved drugs and screening libraries (see Supporting Information). We augmented this set with 39,436 Murcko scaffolds with MW < 300 extracted from Life Chemicals high-throughput screening library (502,902 compounds) for a final set of 39,859 unique ring structures, scaffolds, and chemotypes. Next, we scanned selective ROBIN RNA binders (Gini coefficient > 0.75) and SMM non-RNA binders for existence of these scaffolds and compared the two sets. Table 2 contains the 10 most enriched substructures in selective RNA binders with MW > 100. Diverse nitrogen heterocycles are enriched in RNA-binding compounds, which is consistent with their ability to engage in stacking and hydrogen bonding interactions that have been shown to be critical in small molecule RNA recognition previously. ^[14b]^

**Table 2.**
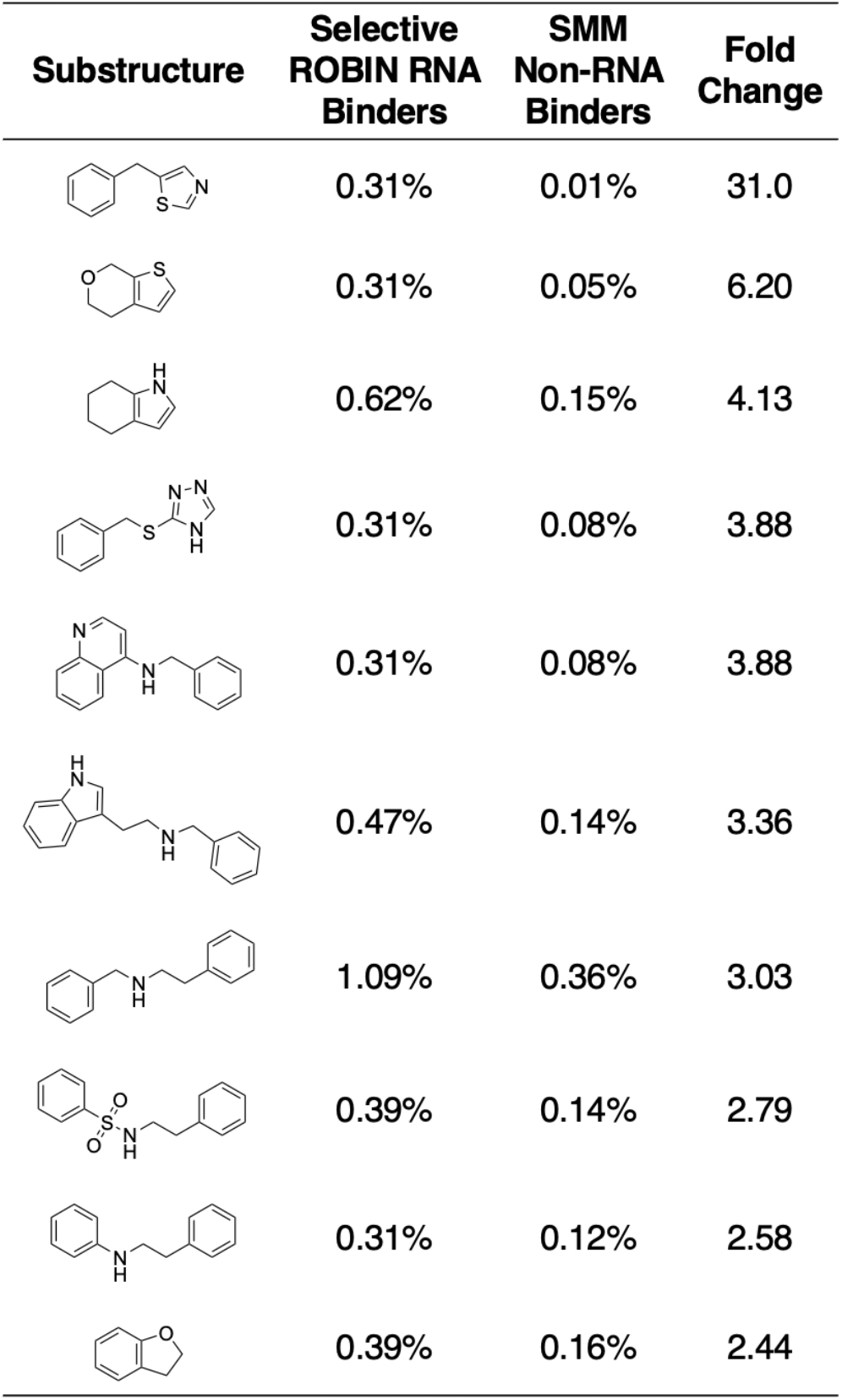
Substructures enriched in selective ROBIN RNA binders. Ten scaffolds and ring structures enriched in selective ROBIN RNA binders relative to SMM Non-RNA binders. Values indicate frequency of occurrence and fold change in binders relative to non-binders.

## Materials and Methods

### Data Preparation

To generate molecular descriptors from our compound libraries, we first obtained their SDF files and generated a single 3D conformer for each compound using DataWarrior’s “Generate Conformers” functionality. We used DataWarrior’s “Systematic, low energy bias” algorithm to minimize energy using the MMFF94s+ forcefield with initial torsions adjusted to “From crystallographic database”. After the generation of 3D conformers, we used ICM-Pro modeling software version 3.9 (MolSoft LLC) to remove salts and explicit hydrogens (Chemistry/Standardize/Remove Salts, Remove Explicit Hydrogens) from each compound. We also standardized groups and tautomers to ensure consistency of our chemical structures (Chemistry/Standardize/Standardize Groups, Standardize Tautomers). Next, we set formal charges for ionizable groups at pH 7.4 consistent with the pH used in our SMM screening buffers (Set Formal Charges/Auto Using pKa Model/pH Value: 7.40). We fed our final standardized 3D SDF files to Mordred for creation of 2D and 3D molecular descriptors. The command-line version of Mordred (version 1.2.0) was used to generate CSV files with 1,827 molecular descriptors for each compound. Descriptors that had missing values in more than 40% of the compounds in each dataset were excluded from analysis. In other cases, missing values were replaced with the median value for that descriptor from the remainder of the dataset. Values approaching infinity were also replaced by the median of their feature column. Full descriptions of chemical descriptors shown in this paper can be found at (https://mordred-descriptor.github.io/documentation/master/descriptors.html).

### Machine Learning

For machine learning models used for prediction, 10% of the compounds (same class balance as the original dataset) were held out as a final holdout/test set. The remaining 90% of the data in each dataset underwent standardization using scikit-learn package’s “StandardScaler” functionality. Model selection and hyperparameter optimization were performed using a stratified 10-fold cross-validation strategy. In class-balanced datasets, mean AUROC across the 10 cross-validation folds was used as the main performance metric. In class-imbalanced cases, mean AUPRC was used as the performance metric. After model selection and hyperparameter optimization, the selected machine learning model was trained on the entire 90% of the data used for cross-validation and tested against the 10% of the data held out in the beginning. The training set in this case (90% of the data) was standardized separately from the test set to prevent any data leakage. Relevant performance metrics on the holdout/test set were reported. The code used for generation of all machine learning models and figures is publicly available on GitHub (https://github.com/ky66/ROBIN).

### Machine Learning Packages

Scikit-learn package (version 1.0.2) within Python (version 3.8.2) was used as the general framework for training of all machine learning models used in the study. Scikit-learn’s “liblinear” solver was used in all cases of LASSO logistic regression optimization. For neural networks, Keras package (version 2.4.3) was used with a Tensorflow backend (version 2.3.1). L2 regularization was applied at each hidden layer and Adam was used as the optimizer with a binary cross-entropy loss function. To interpret the neural networks used, SHAP package was deployed (https://github.com/slundberg/shap) (version 0.37.0) on the Keras models using the deep learning variation of SHAP estimation (Deep SHAP) with 1,000 randomly selected background samples.

### Substructure Searching

ICM-Pro’s “Annotate Chemical by Substructure” tool was used to search each compound for the existence of 39,859 unique ring structures, scaffolds, and chemotypes (the composition of substructures is described in the Results section). For the comparison of selective ROBIN RNA binders and SMM non-RNA binders, we filtered out substructures that occurred in less than 0.25% of selective ROBIN RNA binders.

### TMAP Visualization

TMAP used in this work was created using the Reymond group’s TMAP package in Python (https://github.com/reymond-group/tmap).^[28]^ The embeddings used in the TMAP were created using ECFP4 fingerprints calculated using the RDKit package.

## Discussion

Targeting RNA with small molecules has become a promising area in medicinal chemistry. However, there remain comparatively few examples of small molecules with drug-like physical properties that bind to RNA. The work disclosed here represents a complete account and in-depth analysis of 36 high-throughput screens against diverse nucleic acid structures from therapeutically relevant genes/transcripts. The full composition of a 24,572-compound screening library used to perform these screens is included as Supporting Information. We use machine learning algorithms including logistic regression (LASSO) and neural networks to develop predictive algorithms for RNA binding based on the results of these screens. These algorithms achieve outstanding performance in classifying a library of potential RNA binders, reaching an AUPRC value of 0.78. Evaluation of selected examples of known RNA and protein binders provides further evidence for the model’s utility. Furthermore, analysis of chemical structures found in sets of both RNA binders and protein binders offers insights into the types of chemical matter that bind preferentially to RNA.

It has been shown that RNAs can adopt folds that possess hydrophobic pockets suitable for small molecule binding that are by many metrics similar to hydrophobic pockets on proteins.^[9]^ While examples of small molecules that bind to RNA have been published, it has proven highly challenging to identify compounds that both bind to RNA and have desirable physical properties. Toward this end, the chemical library reported here was not assembled to represent RNA-binding chemical space but rather to represent compounds likely to be suitable as starting points for medicinal chemistry campaigns. Factors considered when designing the library include commercial availability, synthetic tractability, drug-like properties including solubility, cell permeability, and molecular weight, and lack of reactive/promiscuous functional groups (e.g., “PAINS”^[19]^). Also included are a large set of FDA-approved drugs and mechanistically characterized inhibitors. The compounds in this library were screened against diverse RNA and DNA structures, including hairpins, triple helices, three-way junctions, pseudoknots, and G-quadruplexes, providing a broad measure of the ability of small molecules to recognize common nucleic acid structures. Analysis of these screens demonstrates that hit rates for many nucleic acids are comparable to rates frequently seen for proteins, ranging from 0.22% to 1.19% for an individual screen (mean hit rates for all classes was below 1%). When selectivity is factored in hit rates decrease considerably, facilitating the removal of promiscuous/non-specific nucleic acid binders. Here, it is clear that structures such as RNA G4s, pseudoknots, and three-way junctions have the highest selective hit rates. Previous work has indicated that these classes of structure often have suitable hydrophobic pockets and are associated with diverse biological functions, further highlighting their potential as RNA targets.^[37]^ Quantitative assessment of binding selectivity using the Gini coefficient metric revealed that while many molecules were nonselective, the majority exhibited more selective behavior, with roughly two-thirds of hits having a Gini coefficient greater than 0.75.

We analyzed distributions of common drug-like properties such as molecular weight, hydrogen bond donors/acceptors, nitrogen content, total polar surface area, and predicted aqueous solubility (SLogP) for RNA binders identified in ROBIN. These parameters, however, are insufficient to distinguish RNA binders from FDA-approved drugs. We used a set of 1,664 chemical descriptors of each compound to perform a binary classification between FDA-approved drugs and RNA binders from ROBIN with LASSO logistic regression. This algorithm identified a panel of 41 chemical descriptors that was able to differentiate RNA binders from FDA-approved drugs (AUROC = 0.87). Although complex, these parameters contain some commonalities. For example, several features relating to nitrogen content were identified (e.g., SsNH3, SssNH2, SssNH, and AMID_N), indicating the importance of amines and nitrogenous heterocycles in RNA binders. RNA binders were found to have higher values for descriptors describing aromatic ring systems (e.g., n5aHRing, and SaasC), likely due to the propensity of such systems to stack with nucleobases. In contrast, higher values for several oxygen-related descriptors (e.g., NdO, AMID_O, and nAcid) were found in FDA-approved drugs over RNA binders.

Because FDA-approved drugs often contain unusually complex structures such as those derived from natural products, we also assembled a different set of protein binders extracted from the BindingDB database identified from medicinal chemistry programs. We extracted a set of 77,678 compounds with *K*_*d*_ or *K*_*i*_ < 10 nM with a protein target from BindingDB. We visualized the distribution of ROBIN RNA binders, FDA-approved drugs and BindingDB protein binders using their ECFP4 fingerprints on a TMAP to identify clustering patterns of these libraries. This analysis revealed that RNA binders from ROBIN represent diverse scaffolds that are well distributed within the represented chemical space. One potential utility of this TMAP could be identifying FDA-approved drugs or validated inhibitor scaffolds enclosed within RNA-binding branches. For example, in Figure 2, we identified several potential RNA binders similar to ozanimod, a sphingosine-1-phosphate receptor modulator. This information could be useful to identify RNA off-targets of known drugs or to design RNA-targeted molecules from known bioactive scaffolds. The approach of using core RNA-binding scaffolds to identify targets of FDA-approved drugs was recently used to characterize palbociclib as a potent binder of the HIV TAR hairpin.^[38]^ A separate approach also recently indicated that protein-targeting FDA approved drugs similar to known RNA binders can also have RNA-binding character.^[39]^

We also used LASSO logistic regression and neural networks to classify RNA binders from 77,678 protein binders from traditional medicinal chemistry programs (as identified in BindingDB). In this case, we generated an augmented set of RNA binders based on Tanimoto similarity to compounds identified from SMM screens to generate a large dataset that avoids imbalanced classification issues. On this augmented dataset, the LASSO model returned a mean AUPRC of 0.37, while the neural network algorithm performed with an AUPRC of 0.78. Generally, features found to be important for RNA binding in both algorithms included Lipinski parameters, features describing Van der Waals surface area, heteroaromatic ring content, and measures of topological charge. Together, these algorithms identify a complex set of molecular features that can be used to describe and predict RNA-binding behavior within a set of drug-like molecules. Intriguingly, a recent study by Hargrove and co-workers also reported the importance of Van der Waals surface area in RNA-binding behavior in a completely different set of molecules, highlighting the importance of this property.^[14c]^ In addition to performing classification, we also identified chemical scaffolds that are enriched in selective RNA binders (Table 2). In this analysis, the most enriched substructures contain nitrogen heterocycles, including both saturated and unsaturated rings. This finding is consistent with previous reports highlighting the importance of hydrogen bonding and pi stacking for RNA-small molecule interactions.^[14b]^ Benzimidazole, aminoquinoline, and indole structures, frequently seen in reports of RNA binders, were also identified.

This work introduces ROBIN, a new experimentally derived library of 2,188 nucleic acid-binding small molecules, including 2,003 molecules that bind to RNA. To our knowledge, the ROBIN library, and the associated screening dataset, is the largest fully public, experimentally derived investigation of RNA-binding small molecules. The work described here includes information on the targetability of RNA hairpins, DNA and RNA G-quadruplexes, RNA triple helices, RNA pseudoknots, and RNA three-way junctions. This work complements highly valuable existing databases such as R-BIND, INFORNA, and NALDB.^[14e, 15c, 17]^ Distinct from other studies, we report the full results of screens, including hits/binders and non-binders, for a total of 1,627,072 probed binding interactions on SMMs. We used the RNA-binding set of hits to classify RNA binders from FDA-approved drugs and protein-binding ligands using machine learning. While the common drug-like properties of ROBIN RNA binders compare favorably to those of drug-like protein binders, our machine learning analysis reveals molecular descriptors relating to Van der Waals surface area, nitrogen-containing ring systems, and charge differ between the two sets. This dataset will likely find broad use in other informatic applications. As one example, this dataset or the approach described herein could be used to develop screening libraries tailored to identify novel RNA-binders, or ligands specific for classes of structure. This study demonstrates that machine learning-based approaches can be used to characterize RNA-binding chemical space and reveal complex sets of molecular properties that are highly predictive of RNA binding within drug-like chemical libraries. Key to this work was access to a large collection of high throughput screening data on diverse nucleic acid targets, a dataset that is now public. This work is an important step toward developing RNA-targeted small molecules as novel therapeutics.

## Acknowledgments

The authors would like to thank Marc Nicklaus, Ph.D., Megan Peach, Ph.D., and Curran Rhodes, Ph.D. for helpful discussions during the preparation of this manuscript. We thank NCATS and the National Cancer Institute for providing samples of some of the compounds used in this study. This research was supported by the Intramural Research Program of the National Institutes of Health, National Cancer Institute, Center for Cancer Research, project number Z01 BC011585 07 (PI, J. S. Schneekloth, Jr). This work utilized the computational resources of the NIH HPC Biowulf cluster (http://hpc.nih.gov).

## Supporting Information

Code used for machine learning algorithms and tables summarizing the results of SMM screening are available on GitHub (https://github.com/ky66/ROBIN). Supporting datasets, including composition of the SMM screening library, the ROBIN database, the augmented ROBIN dataset, FDA-approved drugs set, BindingDB compounds, and tables of chemical descriptors for these data are available on Figshare (https://doi.org/10.6084/m9.figshare.20401974).

## References

[1] a S. M. Meyer, C. C. Williams, Y. Akahori, T. Tanaka, H. Aikawa, Y. Tong, J. L. Childs-Disney, M. D. Disney, Chem Soc Rev 2020, 49, 7167–7199; b C. M. Connelly, M. H. Moon, J. S. Schneekloth, Jr., Cell Chem Biol 2016, 23, 1077–1090; c A. Umuhire Juru, A. E. Hargrove, J Biol Chem 2021, 296, 100191.

[2] M. J. Hangauer, I. W. Vaughn, M. T. McManus, PLoS Genet 2013, 9, e1003569.

[3] a M. Clamp, B. Fry, M. Kamal, X. Xie, J. Cuff, M. F. Lin, M. Kellis, K. Lindblad-Toh, E. S. Lander, Proc Natl Acad Sci U S A 2007, 104, 19428–19433; b I. Ezkurdia, D. Juan, J. M. Rodriguez, A. Frankish, M. Diekhans, J. Harrow, J. Vazquez, A. Valencia, M. L. Tress, Hum Mol Genet 2014, 23, 5866–5878.

[4] a M. Esteller, Nat Rev Genet 2011, 12, 861–874; b E. Lekka, J. Hall, FEBS Lett 2018, 592, 2884–2900.

[5] a A. L. Hopkins, C. R. Groom, Nat Rev Drug Discov 2002, 1, 727–730; b A. P. Russ, S. Lampel, Drug Discov Today 2005, 10, 1607–1610; c C. V. Dang, E. P. Reddy, K. M. Shokat, L. Soucek, Nat Rev Cancer 2017, 17, 502–508.

[6] a S. T. Crooke, X. H. Liang, B. F. Baker, R. M. Crooke, J Biol Chem 2021, 296, 100416; b T.C. Roberts, R. Langer, M. J. A. Wood, Nat Rev Drug Discov 2020, 19, 673–694.

[7] E. C. Kuijper, A. J. Bergsma, W. Pijnappel, A. Aartsma-Rus, J Inherit Metab Dis 2021, 44, 72–87.

[8] J. R. Thomas, P. J. Hergenrother, Chem Rev 2008, 108, 1171–1224.

[9] a W. M. Hewitt, D. R. Calabrese, J. S. Schneekloth, Jr., Bioorg Med Chem 2019, 27, 2253–2260; b K. D. Warner, C. E. Hajdin, K. M. Weeks, Nat Rev Drug Discov 2018, 17, 547–558.

[10] A. M. Mustoe, S. Busan, G. M. Rice, C. E. Hajdin, B. K. Peterson, V. M. Ruda, N. Kubica, R. Nutiu, J. L. Baryza, K. M. Weeks, Cell 2018, 173, 181–195 e118.

[11] F. A. Abulwerdi, W. Xu, A. A. Ageeli, M. J. Yonkunas, G. Arun, H. Nam, J. S. Schneekloth, Jr., T. K. Dayie, D. Spector, N. Baird, S. F. J. Le Grice, ACS Chem Biol 2019, 14, 223–235.

[12] J. P. Falese, A. Donlic, A. E. Hargrove, Chem Soc Rev 2021, 50, 2224–2243.

[13] a M. M. Attwood, D. Fabbro, A. V. Sokolov, S. Knapp, H. B. Schioth, Nat Rev Drug Discov 2021, 20, 798; b D. Yang, Q. Zhou, V. Labroska, S. Qin, S. Darbalaei, Y. Wu, E. Yuliantie, L. Xie, H. Tao, J. Cheng, Q. Liu, S. Zhao, W. Shui, Y. Jiang, M. W. Wang, Signal Transduct Target Ther 2021, 6, 7; c S. K. Bagal, A. D. Brown, P. J. Cox, K. Omoto, R. M. Owen, D. C. Pryde, B. Sidders, S. E. Skerratt, E. B. Stevens, R. I. Storer, N. A. Swain, J Med Chem 2013, 56, 593–624.

[14] a B. S. Morgan, J. E. Forte, R. N. Culver, Y. Zhang, A. E. Hargrove, Angew Chem Int Ed Engl 2017, 56, 13498–13502; b G. Padroni, N. N. Patwardhan, M. Schapira, A. E. Hargrove, RSC Med Chem 2020, 11, 802–813; c M. Z. Z. Cai, O. Akande, A. Hargrove, ChemRxiv, 2021; d A. Donlic, E. G. Swanson, L.-Y. Chiu, S. L. Wicks, A. U. Juru, Z. Cai, K. Kassam, C. Laudeman, B. G. Sanaba, A. Sugarman, E. Han, B. S. Tolbert, A. E. Hargrove, bioRxiv 2022, 2022.2003.2014.484334; e B. S. Morgan, B. G. Sanaba, A. Donlic, D. B. Karloff, J. E. Forte, Y. Zhang, A. E. Hargrove, ACS Chem Biol 2019, 14, 2691–2700.

[15] a T. Tran, M. D. Disney, Nat Commun 2012, 3, 1125; b H. S. Haniff, L. Knerr, X. Liu, G. Crynen, J. Bostrom, D. Abegg, A. Adibekian, E. Lekah, K. W. Wang, M. D. Cameron, I. Yildirim, M. Lemurell, M. D. Disney, Nat Chem 2020, 12, 952–961; c M. D. Disney, A. M. Winkelsas, S. P. Velagapudi, M. Southern, M. Fallahi, J. L. Childs-Disney, ACS Chem Biol 2016, 11, 1720–1728.

[16] N. F. Rizvi, J. P. Santa Maria, Jr., A. Nahvi, J. Klappenbach, D. J. Klein, P. J. Curran, M. P. Richards, C. Chamberlin, P. Saradjian, J. Burchard, R. Aguilar, J. T. Lee, P. J. Dandliker, G. F. Smith, P. Kutchukian, E. B. Nickbarg, SLAS Discov 2020, 25, 384–396.

[17] S. Kumar Mishra, A. Kumar, Database (Oxford) 2016, 2016.

[18] L. A. Mathews Griner, R. Guha, P. Shinn, R. M. Young, J. M. Keller, D. Liu, I. S. Goldlust, A. Yasgar, C. McKnight, M. B. Boxer, D. Y. Duveau, J. K. Jiang, S. Michael, T. Mierzwa, W. Huang, M. J. Walsh, B. T. Mott, P. Patel, W. Leister, D. J. Maloney, C. A. Leclair, G. Rai, A. Jadhav, B. D. Peyser, C. P. Austin, S. E. Martin, A. Simeonov, M. Ferrer, L. M. Staudt, C. J. Thomas, Proc Natl Acad Sci U S A 2014, 111, 2349–2354.

[19] J. B. Baell, G. A. Holloway, J Med Chem 2010, 53, 2719–2740.

[20] a S. N. Journey, S. L. Alden, W. M. Hewitt, M. L. Peach, M. C. Nicklaus, J. S. Schneekloth, Jr., Medchemcomm 2018, 9, 2000–2007; b D. R. Calabrese, K. Zlotkowski, S. Alden, W. M. Hewitt, C. M. Connelly, R. M. Wilson, S. Gaikwad, L. Chen, R. Guha, C. J. Thomas, B. A. Mock, J. S. Schneekloth, Jr., Nucleic Acids Res 2018, 46, 2722–2732; c C. M. Connelly, T. Numata, R. E. Boer, M. H. Moon, R. S. Sinniah, J. J. Barchi, A. R. Ferre-D’Amare, J. S. Schneekloth, Jr., Nat Commun 2019, 10, 1501; d N. Patel, F. Abulwerdi, F. Fatehi, I. W. Manfield, S. Le Grice, J. S. Schneekloth, Jr., R. Twarock, P. G. Stockley, J Mol Biol 2022, 434, 167557; e M. Yang, S. Carter, S. Parmar, D. D. Bume, D. R. Calabrese, X. Liang, K. Yazdani, M. Xu, Z. Liu, C. J. Thiele, J. S. Schneekloth, Nucleic Acids Res 2021, 49, 7856–7869.

[21] C. M. Connelly, F. A. Abulwerdi, J. S. Schneekloth, Jr., Methods Mol Biol 2017, 1518, 157–175.

[22] A. Ursu, J. L. Childs-Disney, A. J. Angelbello, M. G. Costales, S. M. Meyer, M. D. Disney, ACS Chem Biol 2020, 15, 2031–2040.

[23] D. H. Drewry, C. I. Wells, D. M. Andrews, R. Angell, H. Al-Ali, A. D. Axtman, S. J. Capuzzi, J. M. Elkins, P. Ettmayer, M. Frederiksen, O. Gileadi, N. Gray, A. Hooper, S. Knapp, S. Laufer, U. Luecking, M. Michaelides, S. Muller, E. Muratov, R. A. Denny, K. S. Saikatendu, D. K. Treiber, W. J. Zuercher, T. M. Willson, PLoS One 2017, 12, e0181585.

[24] C. A. Lipinski, F. Lombardo, B. W. Dominy, P. J. Feeney, Adv Drug Deliv Rev 2001, 46, 3–26.

[25] D. F. Veber, S. R. Johnson, H. Y. Cheng, B. R. Smith, K. W. Ward, K. D. Kopple, J Med Chem 2002, 45, 2615–2623.

[26] H. Moriwaki, Y. S. Tian, N. Kawashita, T. Takagi, J Cheminform 2018, 10, 4.

[27] R. Tibshirani, Journal of the Royal Statistical Society: Series B (Methodological) 1996, 58, 267–288.

[28] D. Probst, J. L. Reymond, J Cheminform 2020, 12, 12.

[29] T. Liu, Y. Lin, X. Wen, R. N. Jorissen, M. K. Gilson, Nucleic Acids Res 2007, 35, D198–201.

[30] a D. Rogers, M. Hahn, J Chem Inf Model 2010, 50, 742–754; b S. Riniker, G. A. Landrum, J Cheminform 2013, 5, 26; c M. Awale, J. L. Reymond, J Chem Inf Model 2019, 59, 10–17.

[31] T. Sterling, J. J. Irwin, J Chem Inf Model 2015, 55, 2324–2337.

[32] Y. C. Martin, J. L. Kofron, L. M. Traphagen, J Med Chem 2002, 45, 4350–4358.

[33] S. M. Lundberg, S.-I. Lee, ArXiv 2017, abs/1705.07874.

[34] Y. Ren, Y. F. Wang, J. Zhang, Q. X. Wang, L. Han, M. Mei, C. S. Kang, Clin Epigenetics 2019, 11, 29.

[35] J. Sztuba-Solinska, S. R. Shenoy, P. Gareiss, L. R. Krumpe, S. F. Le Grice, B. R. O’Keefe, J. S. Schneekloth, Jr., J Am Chem Soc 2014, 136, 8402–8410.

[36] J. A. Howe, H. Wang, T. O. Fischmann, C. J. Balibar, L. Xiao, A. M. Galgoci, J. C. Malinverni, T. Mayhood, A. Villafania, A. Nahvi, N. Murgolo, C. M. Barbieri, P. A. Mann, D. Carr, E. Xia, P. Zuck, D. Riley, R. E. Painter, S. S. Walker, B. Sherborne, R. de Jesus, W. Pan, M. A. Plotkin, J. Wu, D. Rindgen, J. Cummings, C. G. Garlisi, R. Zhang, P. R. Sheth, C. J. Gill, H. Tang, T. Roemer, Nature 2015, 526, 672–677.

[37] a D. W. Staple, S. E. Butcher, PLoS Biol 2005, 3, e213; b D. P. Giedroc, P. V. Cornish, Virus Res 2009, 139, 193–208; c P. Kharel, G. Becker, V. Tsvetkov, P. Ivanov, Nucleic Acids Res 2020, 48, 12534–12555.

[38] M. D. Shortridge, V. Vidalala, G. Varani, bioRxiv 2022, 2022.2001.2020.477126.

[39] L. Fang, W. A. Velema, Y. Lee, X. Lu, M. G. Mohsen, A. M. Kietrys, E. T. Kool, bioRxiv 2022, 2022.2007.2018.500496.

