## Supplementary Information for "Machine Learning Informs RNA-Binding Chemical Space"

***Supporting Information***

**Table S1.** Function and associated disease for each nucleic acid target screened.

| **Nucleic Acid Target** | **Biomolecule Type** | **Structure Type** | **Function** | **Associated Disease** |
| --- | --- | --- | --- | --- |
| KRAS | DNA | G-quadruplex | Gene expression | Cancer |
| MTOR | DNA | G-quadruplex | Gene expression | Cancer |
| MYB | DNA | G-quadruplex | Gene expression | Cancer |
| MYC_Pu22 | DNA | G-quadruplex | Gene expression | Cancer |
| VEGF | DNA | G-quadruplex | Gene expression | Cancer |
| RB1 | DNA | G-quadruplex | Gene expression | Cancer |
| BCL2 | DNA | G-quadruplex | Gene expression | Cancer |
| CKIT | DNA | G-quadruplex | Gene expression | Cancer |
| MYCN | DNA | G-quadruplex | Gene expression | Cancer |
| NRAS | RNA | G-quadruplex | Gene expression | Cancer |
| TERRA | RNA | G-quadruplex | Telomere Function | Cancer |
| EWSR1 | RNA | G-quadruplex | Splicing | Cancer |
| AKTIP | RNA | G-quadruplex | Translation | Cancer |
| Zika_NS5 | RNA | G-quadruplex | Viral genome | Zika Infection |
| Zika3PrimeUTR | RNA | G-quadruplex | Viral genome | Zika Infection |
| FGFR | RNA | Hairpin | Splicing | Cancer |
| KLF6_wt | RNA | Hairpin | Splicing | Cancer |
| KLF6_mut | RNA | Hairpin | Splicing | Cancer |
| BCL_XL | RNA | Hairpin | Splicing | Cancer |
| BCL_XL_SS | RNA | Hairpin | Splicing | Cancer |
| RRE2B | RNA | Hairpin | Viral genome | HIV Infection |
| RRE2B_MeA | RNA | Hairpin | Viral genome | HIV Infection |
| Pre_miR_21 | RNA | Hairpin | microRNA | Cancer |
| Pre_miR_17 | RNA | Hairpin | microRNA | Cancer |
| Pre_miR_31 | RNA | Hairpin | microRNA | Cancer |
| HIV_SL3 | RNA | Hairpin | Viral genome | HIV Infection |
| HBV | RNA | Hairpin | Viral genome | HBV Infection |
| Pro_wt | RNA | Hairpin | Splicing | Hutchinson-Gilford Progeria Syndrome |
| Pro_mut | RNA | Hairpin | Splicing | Hutchinson-Gilford Progeria Syndrome |
| PreQ1 | RNA | Pseudoknot | Riboswitch | Bacterial gene expression/infection |
| SAM_ll | RNA | Pseudoknot | Riboswitch | Bacterial gene expression/infection |
| ZTP | RNA | Pseudoknot | Riboswitch | Bacterial gene expression/infection |
| TPP | RNA | Three-way junction | Riboswitch | Bacterial gene expression/infection |
| Glutamine_RS | RNA | Three-way junction | Riboswitch | Bacterial gene expression/infection |
| MALAT1 | RNA | Triple helix | long non-coding RNA | Cancer |
| ENE_A9 | RNA | Triple helix | long non-coding RNA | Cancer |

| **Nucleic Acid Target** | **Biomolecule Type** | **Structure Type** | **Hit Rate** | **Selective Hit Rate** | **Sequence**  **(5' to 3')** |
| --- | --- | --- | --- | --- | --- |
| KRAS | DNA | G-quadruplex | 0.53 | 0.11 | AGGGCGGTGTGGGAAGAGGGAAGAGGGGGAGGCAG |
| MTOR | DNA | G-quadruplex | 0.31 | 0.09 | GGGGAAGGCGGGCGGTGGGGCAGGGGG |
| MYB | DNA | G-quadruplex | 0.34 | 0.04 | GGAGGAGGAGGTCACGGAGGAGGAGGAGAAGGAGGAGGAGGA |
| MYC_Pu22 | DNA | G-quadruplex | 0.27 | 0.05 | AGGGTGGGGAGGGTGGGG |
| VEGF | DNA | G-quadruplex | 0.48 | 0.08 | CGGGGCGGGCCGGGGGCGGGGT |
| RB1 | DNA | G-quadruplex | 0.49 | 0.07 | CGGGGGGTTTTGGGCGGC |
| BCL2 | DNA | G-quadruplex | 0.45 | 0.06 | AGGGGCGGGCGCGGGAGGAAGGGGGCGGGA |
| CKIT | DNA | G-quadruplex | 0.34 | 0.04 | AGGGAGGGCGCTGGGAGGAGGG |
| MYCN | DNA | G-quadruplex | 0.85 | 0.52 | AGGGGGTGGGAGGGGGCATGCAGATGCAGGGGGT |
| NRAS | RNA | G-quadruplex | 0.5 | 0.08 | UGUGGGAGGGGCGGGUCUGGG |
| TERRA | RNA | G-quadruplex | 0.54 | 0.05 | GGGUUAGGGU |
| EWSR1 | RNA | G-quadruplex | 1.19 | 0.46 | GGGGCAGGGGAAGAGGGGG |
| AKTIP | RNA | G-quadruplex | 0.91 | 0.2 | GGGGUGGGGCGGGGCGGGG |
| Zika_NS5 | RNA | G-quadruplex | 0.7 | 0.23 | GUGGAGGUGGGACGGGAGAGACUCUGGGAGA |
| Zika3PrimeUTR | RNA | G-quadruplex | 0.82 | 0.3 | GCGGCGGCCGGUGUGGGGAA |
| FGFR | RNA | Hairpin | 0.42 | 0.15 | GCUCUUGCUUCGCUUGUUUUCUAGGCCGCCGGUGUUAACACCACGGACAAAGAGC |
| KLF6_wt | RNA | Hairpin | 0.3 | 0.07 | GCUGGCAAUCACGUGCCAGC |
| KLF6_mut | RNA | Hairpin | 0.37 | 0.11 | GCUGGCAAUCACAUGCCAGC |
| BCL_XL | RNA | Hairpin | 0.36 | 0.05 | UUGGAUCCAGGAGAACGGCGGCU |
| BCL_XL_SS | RNA | Hairpin | 0.9 | 0.31 | AGCCUUGGAUCCAGGAGAACGGCGGCUGGGUAAGAACCAAGCCCUUGUGUGUCCCUUUUCUUUGGCCUC |
| RRE2B | RNA | Hairpin | 0.82 | 0.31 | UGGGCGCAGUGUCAUUGACGCUGACGGUACA |
| RRE2B_MeA | RNA | Hairpin | 0.96 | 0.39 | UGGGCGCAGUGUCAUUG(m6A)CGCUG(m6A)CGGUACA |
| Pre_miR_21 | RNA | Hairpin | 0.34 | 0.1 | GGGUUGACUGUUGAAUCUCAUGGCAACCC |
| Pre_miR_17 | RNA | Hairpin | 0.33 | 0.11 | CAAAGUGCUUACAGUGCAGGUAGUGAUAUGUGCAUCUACUGCAGUGAAGGCACUUGUAG |
| Pre_miR_31 | RNA | Hairpin | 0.36 | 0.08 | AGGCAAGAUGCUGGCAUAGCUGUUGAACUGGGAACCUGCUAUGCCAACAUAUUGCCAUC |
| HIV_SL3 | RNA | Hairpin | 0.72 | 0.31 | GACUAGCGGAGGCUAGAA |
| HBV | RNA | Hairpin | 0.22 | 0.07 | GGUUCAUGUCCUACUGUUCAAGCCUCCAAGCUGUGCCUUGGGUGGCUUUGGGGCAUGGACC |
| Pro_wt | RNA | Hairpin | 0.3 | 0.14 | GAGCCCAGGUGGGCGGACCCAUCUCCUCUGGCUC |
| Pro_mut | RNA | Hairpin | 0.25 | 0.08 | GAGCCCAGGUGGGUGGACCCAUCUCCUCUGGCUC |
| PreQ1 | RNA | Pseudoknot | 0.68 | 0.25 | AGAGGUUCUAGCUACACCCUCUAUAAAAAACUAA |
| SAM_ll | RNA | Pseudoknot | 0.95 | 0.35 | UCGCGCUGAUUUAACCGUAUUGCAAGCGCGUGAUAAAUGUAGCUAAAAAGGG |
| ZTP | RNA | Pseudoknot | 0.91 | 0.39 | UAUCAGUUAUAUGACUGACGGAACGUGGAAUUAACCACAUGAAGUAUAACGAUGACAAUGCCGACCGUCUGGGCGAACA |
| TPP | RNA | Three-way junction | 0.66 | 0.11 | CAGUACUCGGGGUGCCCUUCUGCGUGAAGGCUGAGAAAUACCCGUAUCACCUGAUCUGGAUAAUGCCAGCGUAGGGAAGUGCUG |
| Glutamine_RS | RNA | Three-way junction | 0.91 | 0.35 | UAAUCGUUGGCCCAGUUUAUCUGGGUGGAAGUAAGGUCUUUGGCCUGAAGCAACGCGCCUC |
| MALAT1 | RNA | Triple helix | 0.78 | 0.22 | AAAGGUUUUUCUUUUCCUGAGAAAUUUCUCAGGUUUUGCUUUUUAAAAAAAAAGCAAAA |
| ENE_A9 | RNA | Triple helix | 0.45 | 0.19 | AAAAAAAAA and GGCUGGGUUUUUCCUUCGAAAGAAGGUUUUUAUCCCAGUC |

**Table S2.** Summary of SMM screening of each nucleic acid target.

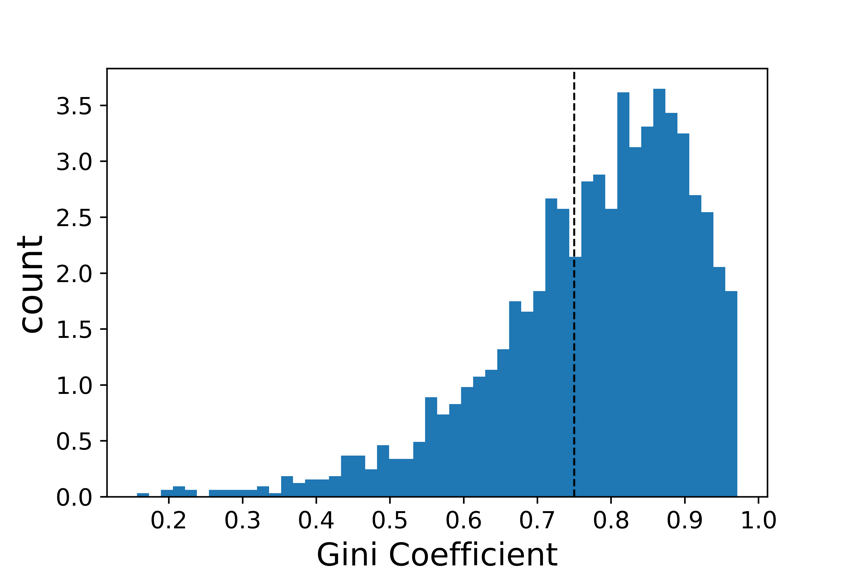

**Figure S1.** Histogram of Gini coefficients calculated for 2,003 compounds

compiled in the ROBIN RNA binding library.
